# Spatially distinct microglia coordinate the response to photoreceptor injury in zebrafish

**DOI:** 10.64898/2026.09.04.749450

**Authors:** Mikiko Nagashima, Lu Jiang, Sriram Garapati, Thanh Hoang, Peter F. Hitchcock

## Abstract

Microglia exhibit substantial molecular and functional heterogeneity, however, how the local tissue environment influences their responses to neuronal injury remains poorly understood. The zebrafish retina provides a unique opportunity to examine this relationship, because in addition to microglia within the retinal parenchyma, it contains a distinct population that resides within the subretinal space. In this study, we characterize the anatomical distribution and molecular heterogeneity of retinal microglia in zebrafish and examine their response to selective photoreceptor injury. Retinal microglia occupied distinct anatomical niches and exhibited diverse transcriptional states, including complement associated, proliferative, and apoc1/apoeb-enriched populations, suggesting that microglial heterogeneity reflects both tissue context and cellular. Following damage both subretinal and parenchymal microglia rapidly alter their morphology and migrate toward the injured site, resulting in their accumulation in the subretinal space. Microglia subsequently proliferate predominantly within this compartment. By 14 days post lesion, when photoreceptor regeneration is nearly completed, microglial morphology, distribution, and the number largely return to their unlesioned state. In contrast to the dynamic response of retinal microglia, response of microglia within the optic tectum was negligible. To investigate the mechanisms underlying the maintenance of these distinct microglial populations and their injury-induced responses, we examined retinal microglia in *csf1ra* mutants. In mutants lacking colony stimulating factor 1 receptor a (Csf1ra), parenchymal microglia are markedly reduced, whereas the number of subretinal microglia is preserved. Photoreceptor death following photolytic injury was comparable between *csf1ra* mutants and wildtype animals, and both parenchymal and subretinal microglia responded to the injury. However, proliferation of microglia in csf1ra mutants was severely impaired, and the normal redistribution of parenchymal microglia was not restored. These findings identify distinct anatomical and molecular features of retinal microglia and demonstrate that Csf1ra signaling differentially regulates microglial maintenance and injury-induced dynamics during photoreceptor regeneration.

## 1 Introduction

Microglia are the innate immune cells of the central nervous system. Following injury, microglia become activated and respond by releasing cytokines and other inflammatory mediators, phagocytosing cellular debris, and aiding the restoration of tissue homeostasis. Although activation of microglia is a common feature of CNS injury, increasing evidence indicates that this complex response is heterogeneous and sharped by the local tissue environment(Ziebell et al. 2015; Ransohoff 2016; Eggen and Kooistra 2026). Recent studies have revealed substantial heterogeneity among microglia, with microglial states exhibiting different transcriptional profiles and functional properties and occupying specialized anatomical niches(O’Koren et al. 2019; Fumagalli et al. 2025). However, it remains unclear to what extent the local tissue environments shape microglial identity and influences how distinct microglial populations respond to neuronal injury.

In the mammalian retina, microglia reside primarily in the inner and outer plexiform layers and the retinal ganglion cell layer(Provis et al. 1996; Santos et al. 2008). In contrast, microglia are normally absent from the outer nuclear layer and the subretinal space, the extracellular compartment between the photoreceptor outer segments and the apical processes of the retinal pigment epithelium (RPE)(Santos et al. 2008). Following photoreceptor damage or degeneration, however, microglia migrate into the outer retina and subretinal space, where they contribute to both tissue repair and, in chronic degenerations, disease pathology(Combadière et al. 2007; Luhmann et al. 2009; Chinnery et al. 2012; Eandi et al. 2016). In zebrafish, in contrast to the mammals, microglia are present not only within the neural retinal parenchyma but also within the subretinal space, where they reside among the somata of the RPE(Chen et al. 2022; Song et al. 2024). The presence of microglia in these anatomically distinct retinal niches raises the possibility that their local environments contribute to differences in their molecular identity and functional properties.

In the zebrafish retina, microglia have been extensively studied in the context of neuronal and photoreceptor regeneration(Iribarne 2021; Nagashima and Hitchcock 2021; Mitchell et al. 2018, 2019; Song et al. 2024). In zebrafish, retinal regeneration involves the reprogramming of Müller glia and subsequent proliferation and differentiation of Müller glia-derived progenitors(Lenkowski and Raymond 2014; Nagashima et al. 2013; Lahne et al. 2020; Hoang et al. 2020). Recent studies have demonstrated that acute microglia-mediated inflammatory response governs the initiation and regulation of this regenerative response(White et al. 2017; Silva et al. 2020a; Iribarne and Hyde 2022; Bludau et al. 2024). We, and others, have also reported that failure to resolve inflammation can compromise survival and maturation of regenerating photoreceptors(Silva et al. 2020b; Magner et al. 2022). These findings highlight the importance of tightly regulated microglial responses during retinal regeneration.

Despite the established importance of microglia in retinal regeneration, it remains unclear how microglial molecular heterogeneity relates to their anatomical distribution and response to injury. The presence of microglia in both retinal parenchyma and subretinal space, together with evidence for molecularly distinct microglial populations, raises the possibility that microglial responses to photoreceptor injury may differ according to their cellular state and tissue context. However, it remains unknown whether distinct microglial populations exhibit different injury responses or instead converge on a common regenerative response. Determining how anatomical niche influences microglial responses to injury is important for understanding how microglial heterogeneity and tissue context shape inflammation during retinal regeneration.

## 2 Materials and methods

### 2.1. Animals

Wildtype (AB-strain, Danio rerio; ZIRC, University of Oregon, Eugene, OR), transgenic reporter lines, Tg(mpeg1.1:eGFP)^gl22tg^(Ellett et al. 2011a), Tg(mpeg1.1:mCherry)^gl23tg^(Ellett et al. 2011a), and csf1ra^j4e1^ mutant(Parichy et al. 2000) were maintained at 28ºC on a 14/10 hours light/dark cycle, employing standard husbandry protocols. Adult fish between 6 to 12 months of age and of either sex were used. All animal procedures were approved by the Institutional Animal Care and Use Committee at the University of Michigan.

### 2.2. Genotyping

Genotyping for csf1ra^j4e1^ was validated by Sanger sequencing. In brief, genomic DNA was extracted from the clipped tail using hot Alkaline lysis buffer. Genomic DNA regions containing csf1ra^j4e1^ were amplified using PCR primers, (F, 5’-ACT CTT GGT GCT GGT GCG TTT G-3’; R, 5’-CTT TGA GCA TTT TCA CAG CC-3’, (Parichy et al. 2000)) and Phusion High-Fidelity DNA Polymerase (New England BioLabs). To identify Tg(mpeg1.1:mCherry) carriers in the csf1ra^j4e1^ background, standard PCR assay was performed using primer against mCherry construct (F, 5’-GGG CGA GGA GGA TAA CAT GG-3’; R, 5’-GGT GTA GTC CTC GTT GTG GG-3’).

### 2.3. Light lesions

To selectively kill photoreceptors, high intensity light lesions were used(Bernardos et al. 2007; S. Taylor et al. 2012). In brief, 4-6 zebrafish were exposed to 100,000 lux light from an EXFO-X Cite 120W metal halide lamp for 30 min. Following exposure fish were immediately returned to the recirculating habitats and normal light/dark cycles. Survival times are established from light onset to when animals are sacrificed.

### 2.4. Immunohistochemistry on retinal cross section

Eyes were fixed in 4% paraformaldehyde with 5% sucrose in 0.1M phosphate buffer, pH 7.4, at 4C overnight. After rinsing with 5% sucrose in phosphate buffer, eyes were cryoprotected via increasing concentrations of sucrose, embedded in OCT-20% sucrose solution, and sectioned at 6 µm. Immunocytochemistry was performed as previously described(Nagashima et al. 2020). In brief, non-specific antibody binding was blocked with PBS containing 20% normal goat serum, 0.5% Triton X-100, and 0.1% sodium azide. Primary antibodies (mouse anti-4C4(Rovira et al. 2023) 1:500; rabbit anti-PCNA Abcam ab18197, 1:400: chicken anti-GFP, Abcam ab13970, 1:500) were diluted with PBS containing 1% normal goat serum, 0.5% Triton X-100, and 0.1% sodium azide. Incubation was performed overnight at 4ºC. After washing, slides were incubated with secondary antibodies (Alexa Fluor 488 goat anti-chicken IgG, 1:200; Alexa Fluor 555 goat anti-mouse IgG, 1:200; Alexa Fluor 647 goat anti-rabbit IgG, 1:200) at room temperature for 2 hrs. After washing, sections were stained with Hoechst 33342 (Thermo Fisher Scientific) and coverslipped with ProLong Gold (Thermo Fisher Scientific). For immunostaining against PCNA, slides were pretreated with sodium citrate buffer at 95ºC for 5 min before the blocking step.

### 2.4. Flat-mount retinal immunocytochemistry

To perform immunocytochemistry in retinal-RPE preparations, we slightly modified the previously published protocol(Nagashima et al. 2017; Nagashima and Hitchcock 2020). In brief, retinal-RPE tissue was carefully dissected from a fixed globe. A large ventral cut was made to establish the orientation of the dissected retinas. To prepare tectal sample, tectul tissues were carefully dissected from fixed whole-head preparation. After rinsing with 5% sucrose in phosphate buffer, pH7.4, melanin in the RPE was bleached in 20% hydrogen peroxide solution for 10-20 min at 55ºC. Isolated retina/RPE were then washed with PBS/0.5% triton-X for 10 min at room temperature before blocking non-specific binding by antibodies in PBS containing 10% normal goat serum, 1% Tween 20, 1% Triton-X, 1% DMSO, and 0.5% sodium azide. Primary antibodies (rabbit anti-GFP Invitrogen A11122, 1:200; mouse anti-ZO-1 (ZO1-1A12), Invitrogen, 33-9100 1:200) and secondary antibodies (Alexa Fluor 555 goat ant-mouse IgG, 1:200; Alexa Fluor 647 goat anti-rabbit IgG, 1:200) were diluted in PBS containing 0.5% normal goat serum, 1% Tween 20, 1% Triton-X, 1% DMSO and 0.5% sodium azide. Retina/RPE preparations were mounted, RPE side down, on microscope slides and coverslipped using Prolong Gold.

### 2.5. Microscope and Image analysis

Retinal cross sections were imaged with DM6000 Upright Microscope System (Leica Microsystems). Flat-mount preparations were imaged using a Leica SP5 confocal microscope (Leica Microsystems) or a Leica STELLARIS 8 FALCON confocal microscope (Leica Microsystems). Application Suite X (Leica Microsystems) and were used for 3D reconstruction.

### 2.6. Cell counts and statistical analyses

Image analyses were performed using Application Suite X (Leica Microsystems) and Image J (https://imagej.nih.gov/ij/). For flat mount retinas, microglia were counted in a total average area of 167,682 mm2/retina from 3 to 5 individual animals. In each quadrant of each retina, areas lying between the optic disc and margin were sampled. Individual layers were identified relative to the boundaries demarcated by ZO1-labeled adherens and tight junctions at the outer limiting membrane and between RPE cells, respectively. In sections, PCNA-labeled cells were counted along an average of 320.43 mm length of each section. Statistical analysis was performed using a student’s t-test (GraphPad Prism Software). p-values ≤ were considered to indicate statistically significant differences.

### 2.7. Single-cell RNA-sequencing analysis of microglial subtypes

Microglia/macrophage profiles were extracted from a previously annotated zebrafish retinal single-cell RNA-sequencing dataset(Lyu et al. 2025; Nagashima et al. 2026). Cells annotated as Microglia_ImmuneCells from control (unlesioned) retinas were selected, yielding 2,335 cells. The log-normalized expression matrix stored in the .raw layer of the AnnData object was used for downstream analysis and marker visualization.

Genes detected in fewer than five cells were excluded before dimensionality reduction. Up to 3,000 highly variable genes were selected using the Seurat dispersion-based method implemented in Scanpy. Mitochondrial and ribosomal genes were excluded from the highly variable gene set but retained for quality-control and marker analyses. Expression values were scaled with a maximum value of 10, followed by principal-component analysis using 30 components. Sample-associated variation was corrected using Harmony with library identifier as the batch variable. A 20-nearest-neighbor graph was constructed from the Harmony-corrected principal components, and UMAP coordinates were calculated with min_dist = 0.3.

Microglial subclusters were identified by Leiden clustering at a resolution of 0.2 using an undirected graph and a random seed of 0.

The source dataset had undergone doublet removal before subclustering. Residual doublets were independently evaluated in each control library using Scrublet and DoubletDetection. Because the original feature-barcode matrices were unavailable, integer counts were reconstructed from the stored log-normalized expression matrix and total-count metadata. Cells identified as doublets by both methods were considered high-confidence residual doublets and excluded. Cell identities were further assessed based on cluster-enriched genes, library complexity, canonical myeloid markers, and retinal lineage markers. Clusters exhibiting coordinated neuronal, photoreceptor, other retinal glial, or low-complexity ribosomal expression programs were excluded. The analysis workflow was repeated following major filtering steps. Following filtering, 651 microglia-enriched cells remained.

Cluster-enriched genes were identified by Wilcoxon rank-sum testing using log-normalized expression. For visualization, dot size represents the fraction of cells expressing each gene and color represents scaled mean expression. Violin plots show log-normalized expression distributions within each cluster, with widths normalized within clusters and therefore not representing absolute cell numbers. Analyses were performed in Python 3.11 using Scanpy 1.10.4, AnnData 0.11.3, harmonypy 2.0.0, and DoubletDetection 4.3.0.post1.

## 3. Results

### 3.1. Microglia in the zebrafish retina exhibits positional and transcriptional diversity

For zebrafish, antibodies and microglia-specific fluorescent reporter lines are routinely used to visualize microglia(Ellett et al. 2011b; Becker and Becker 2001). The monoclonal antibody, 4C4, recognizes Galectin-3 binding protein and is specific to microglia(Rovira et al. 2023). In the retinal parenchyma of adult zebrafish, 4C4-labels microglia that form thin tangential sheets within the retinal ganglion cell layer/nerve fiber layer, at the interface of the inner plexiform and nuclear layers, and within the thin outer plexiform layer (Fig. 1A,B).

**Figure 1.**
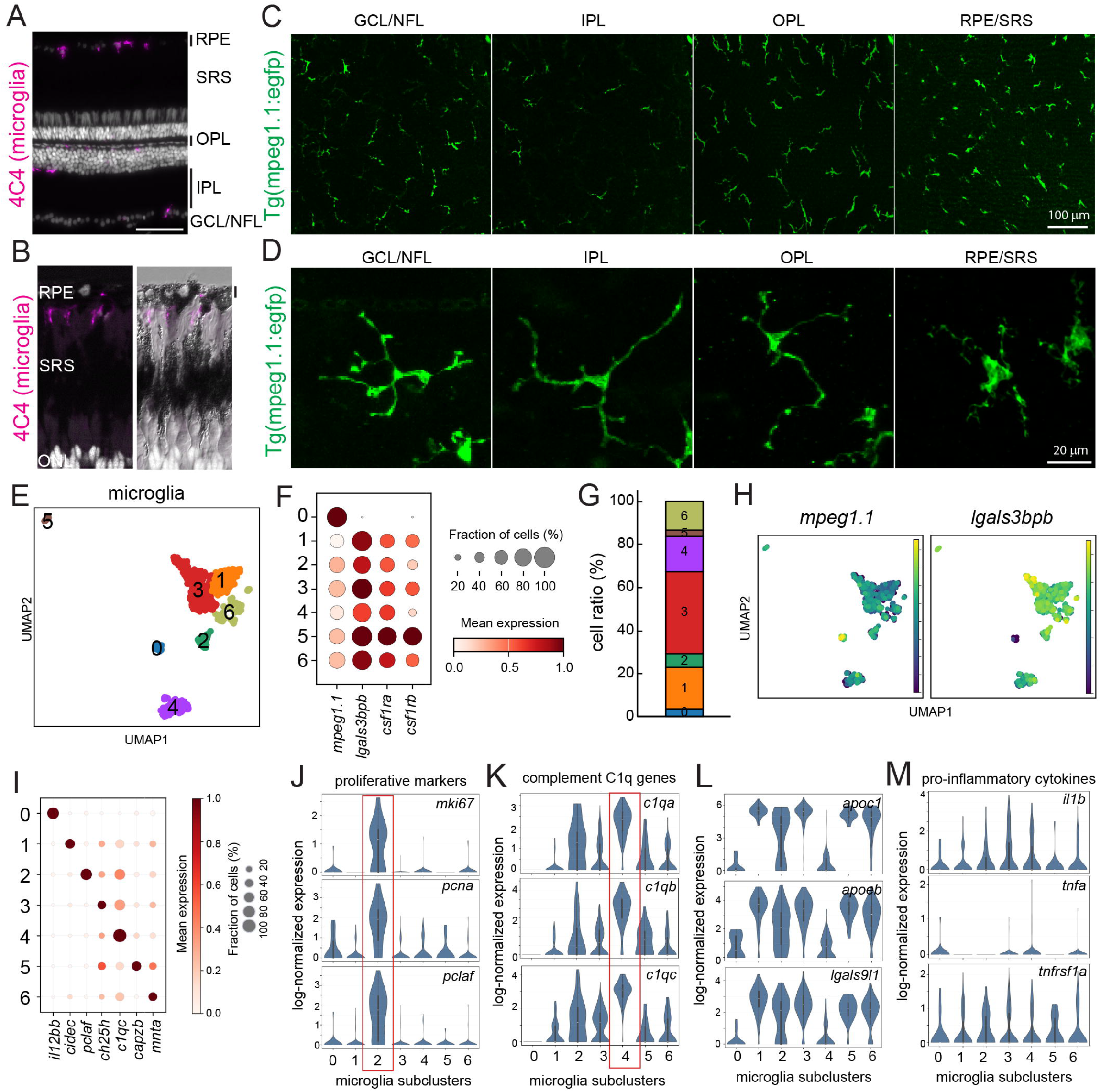
Spatial and transcriptional heterogeneity of microglia in adult zebrafish retina. (A,B) Cross section of unlesioned retina stained with microglia specific antibody, 4C4. (C) Maximum projection of Z-stack series of flat-mount retinal preparation. Tg(mpeg1.1:eGFP)-labeled microglia at the different retinal layers. (D) High magnification image of eGFP+ microglia in different retinal layers. (E) UMAP plot of microglial populations from unlesioned retina. (F) Dot plot of selected canonical microglial markers. (G) Fraction of cells in each clusters. (H) mRNA expression of mpeg1.1 and lgals3bpb. (I) Dot plot of selected genes uniquely expressed by each clusters. (J-M) Violin plots of proliferative marker (J), complement Cq1 genes (K), apoc1/apoeb/lgals9l (L), and pro-inflammatory cytokines. Scale bars: A 50um; C 100um; D 20um. RPE, retinal pigment epithelium; SRS, subretinal space; ONL outer nuclear layer; OPL outer plexiform layer; INL, inner nuclear layer; GCL/NFL, ganglion cell layer/nerve fiber layer.

Individual parenchymal microglia occupy distinct non-overlapping territories, forming a regularly spaced tiling pattern (Fig. 1C). These microglia have ramified morphologies with small soma, and long, thin processes that radiate from the cell body (Fig. 1D). In addition, and unlike microglia in the mammalian retina, there is a population of microglia in zebrafish that reside among the somata of the retinal pigmented epithelium (Fig. 1A,B also see(Chen et al. 2022; Song et al. 2024)). These subretinal microglia also form a regularly spaced, tangential mosaic (Fig. 1C), though subretinal microglia have slightly larger cell bodies and shorter processes (Fig. 1D). The planimetric density of subretinal microglia is greater than any individual layer of parenchymal microglia (GCL/NFL, 278.8±46.6 cells; IPL, 140.1±46.3 cells; OPL, 344.4±70.2 cells; SRS, 442.9±43.8 cells, per 1mm^2^), representing nearly 35% of all microglia in the zebrafish retina. These results indicate that microglia in adult zebrafish retina are distributed across anatomically distinct niches, with a substantial proportion residing within the subretinal space.

To determine whether retinal microglia also exhibit molecular heterogeneity, we analyzed previously generated single-cell RNA sequencing (scRNAseq) datasets from unlesioned adult zebrafish retina(Lyu et al. 2025; Nagashima et al. 2026). Because the samples were not collected separately from the parenchymal and subretinal populations, these data cannot be used to directly assign transcriptional populations to specific anatomical niches. After filtering and quality control, we retained 651 mpeg1.1+ myeloid cells. Subclustering identified 6 transcriptionally distinct populations with distinct gene expression profiles (Fig. 1E-H). Cluster 0 was distinguished from the other populations by its low or absent expression of *lgals3bpb, csf1ra*, and *csf1rb* (Fig.1F,G, H). Because these markers were consistently expressed in the remaining clusters but largely absent from Cluster 0, we could not confidently assign this population to microglia. We therefore excluded Cluster 0 from subsequent analyses of microglial transcriptional states rather than interpret it as a microglial state. Cluster 2 expressed high levels of proliferative markers, including *mki67, pcna*, and *pclaf*, consistent with a proliferative state (Fig. 1J). Cluster 4 was enriched for *cq1a, cq1b*, and *cq1c*, indicating a complement-associated/homeostatic microglial population (Fig. 1K). Consistent with a previous scRNAseq study of microglia following genetic photoreceptor damage(Weimar et al. 2026), expression of complement C1q genes was inversely correlated with *apoc1* and *apoeb* (Fig. 1L). The *apoc1*/*apoeb*-enriched populations (clusters 1, 2, 3, 5, and 6) also expressed genes associated with microglial activation or stress, including *lgals9l1* (Fig. 1L).

However, these populations did not show substantial expression of canonical pro-inflammatory cytokines, including *il1b, tnfa*, and their associated signaling components (Fig. 1M). Because the retinas were dissociated and processed before single cell capture, these transcriptional profiles may reflect activation or stress induced during tissue preparation rather than stable microglial states present in the intact retina. Thus, scRNAseq reveals substantial transcriptional heterogeneity among retinal microglia-enriched cells in the unlesioned retinas. However, the transcriptomic data did not identify a population that could be confidently assigned to the anatomically distinct subretinal microglia. We therefore used the anatomical distribution and injury response analysis described below to examine whether microglia occupying different retinal niches exhibit distinct dynamics following photoreceptor injury.

### 3.2. Rapid and dynamic migration of microglia in response to photoreceptor injury

To qualitatively and quantitatively analyze parenchymal and subretinal microglia, we developed a novel procedure to isolate, stain, and image eGFP-labeled microglia in flat mount preparations from the Tg(mpeg1.1:eGFP) line (see Materials and Methods). To precisely locate microglia within the retinal layers, retinas were stained with the antibody Zonula Occuludens-1 (ZO1), which labels tight junctions between the somata of the RPE and adherens junctions at the outer limiting membrane(Nagashima et al. 2017). A z-stack series of eGFP-labeled microglia in unlesioned retina reveals the easily discernible layers of parenchymal and subretinal microglia (Fig. 2A,B and Movie 1). To characterize the chemotactic response of microglia following photoreceptor injury, photolytic lesions were made as described in Methods. This lesioning paradigm causes the selective death of photoreceptors in a horizontal band spanning the nasotemporal axis of the retina(Qin et al. 2009). Dorsal to this horizontal band, UV cones are also ablated, whereas ventral to this band of ablated photoreceptors, neither rods nor cones are killed(Nagashima et al. 2013). These patterns of photoreceptor injury allow us to evaluate and contrast the responses of microglia proximal to dying photoreceptors and the responses of microglia proximal to uninjured photoreceptors. Between 1 and 4 hours post lesion (hpl), parenchymal microglia become progressively less abundant, especially within the horizontal band of injured photoreceptors (Fig. 2A). In cross sections, thin, radially oriented processes of microglia span the inner and outer nuclear layers (Fig. 2C), suggesting radial migration of microglia from the retinal parenchyma toward the subretinal space. Simultaneously, between 1 and 4 hpl subretinal microglia adopt elongated, rod-shaped morphologies with sausage-shaped cell bodies and long, thin processes (Figs. 2A,B(Taylor et al. 2014; Vidal-Itriago et al. 2022)). At 4 hpl, the horizontal band of injured photoreceptors is demarcated by a marked accumulation of microglia at the dorsal and ventral boundaries of the lesion (Fig. 2A). Additionally, there are also fewer subretinal microglia in ventral retina (Fig. 2A), suggesting tangential migration of these microglia toward the injured photoreceptors. Microglial counts show that at 4 hpl the average number of parenchymal microglia is significantly reduced in all retinal quadrants, while the average number of subretinal microglia increases dramatically, except for within ventral retina (Fig.2D,E). By 24 hpl, the number of parenchymal microglia overlying the photoreceptor injury is dramatically reduced (Fig. 2G). Together, these results indicate that following photoreceptor injury both parenchymal and sub-retinal microglia migrate rapidly into the subretinal space. Additionally, there is marked tangential migration of subretinal microglia from regions outside the immediate area of photoreceptor injury. The reduction in parenchymal microglia was observed across all retinal quadrants, including regions outside the photoreceptor lesion, indicating that microglial redistribution was not restricted to areas immediately adjacent to dying photoreceptors. These results indicate that photoreceptor injury induces a rapid, coordinated redistribution of microglia throughout the retina. Both parenchymal and subretinal microglia migrate toward the subretinal space, while subretinal microglia also undergo tangential migration from regions outside the immediate area of photoreceptor injury.

**Figure 2.**
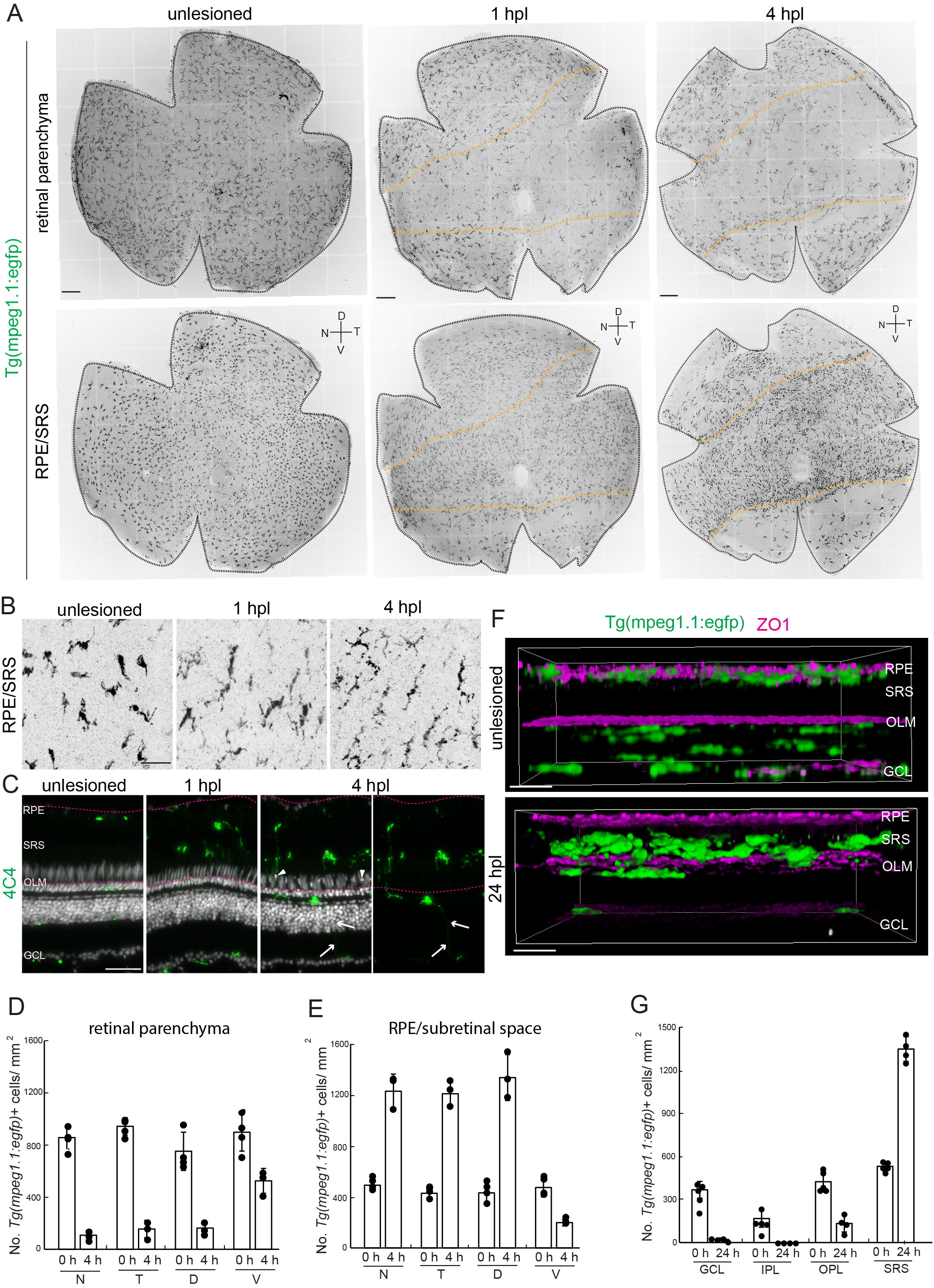
Rapid migration of subretinal and parenchymal microglia following photolytic lesion. (A) flat-mount retinal preparation of Tg(mpeg1.1:eGFP)-labeled microglia within the retinal parenchyma (top) and the subretinal space (bottom) microglia in unlesioned and at 1 and 4 hours post lesion. Dashed lines indicate the boundary of horizontal bands with photoreceptor injury. (B) High magnification image of microglia in the subretinal space in unlesioned and at 1 and 4 hours post lesioned retina. (C) Retinal cross sections stained with 4C4 in unlesioned and 1 and 4 hours post lesioned retinas. Arrows indicate microglial process extending radially. (D,E) Quantification of the number of microglia in retinal parenchyma (D) and subretinal space (E). (F) Orthogonal 3D rendering of retinal flat-mount Z-stack series of Tg(mpeg1.1:eGFP) retina immunostained for ZO1 in unlesioned (top) and 24 hours post lesion (bottom). (G) Quantification of the number of microglia in each retinal layer of the temporal retina. Scale bars: A 200um; B; C; F. RPE, retinal pigment epithelium; SRS, subretinal space; ONL outer nuclear layer; OPL outer plexiform layer; INL, inner nuclear layer; GCL/NFL, ganglion cell layer/nerve fiber layer.

### 3.3. Injury to the photoreceptor does not trigger activation of microglia in the optic tectum

The optic tectum is the primary target of retinal ganglion cell axons in zebrafish and is functionally analogous to the superior colliculus in mammals. It is a highly organized, laminated structure, with the superficial stratum opticum (SO) serving as a major retinorecipient layer (Fig. 3A;(Robles et al. 2013)). Because photoreceptor injury induces a rapid and extensive redistribution of microglia within the retina, we asked whether this response extends to the central visual target of retinal neurons. We first characterized the distribution and morphology of microglia in the optic tectum using cross section stained with 4C4 and flat-mount preparation of the tectum from Tg(mpeg1.1:eGFP). Microglia were readily detected throughout the tectal layers (Fig. 3B). In tectal flat-mount preparations, we focused on microglia located near the superficial surface of the tectum, including the stratum marginale and the SO. This region was selected because the SO is a major retinorecipient layer and therefore provides an appropriate site to assess whether the microglial response to retinal photoreceptor injury extends to the downstream visual circuitry. We found that microglia in these layers frequently displayed elongated cell bodies and processes oriented approximately parallel to the trajectory of the optic nerve (Fig. 3C). This organization was particularly evident at the level of the SO. These observations reveal a close spatial association between tectal microglia and the incoming retinal axons. We next examined whether microglia in the optic tectum respond to photoreceptor injury. At 24 hpl, the morphology, number, and distribution of microglia were qualitatively similar to those observed in unlesioned animals (Fig. 3D), indicating that acute microglial response to selective photoreceptor injury is spatially restricted to the retina and does not extend detectably to the optic tectum. Because the response to tectal microglia was negligible, subsequent analysis focused on retinal microglia.

**Figure 3.**
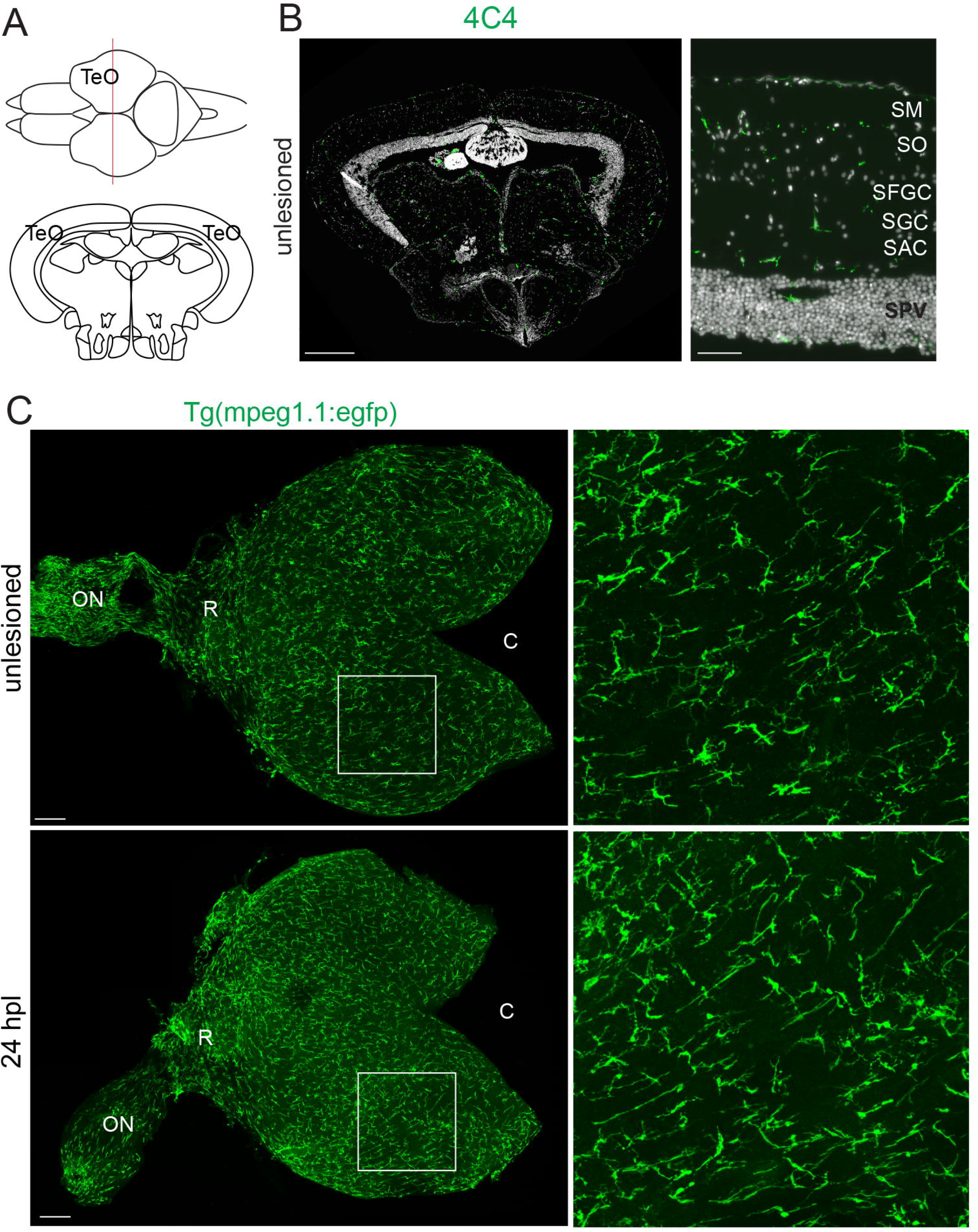
Photoreceptor injury does not activate microglia in the optic tectum. (A) Schematic of optic tectum. (B) Cross section of optic tectum immunostained for 4C4. (C)Flat-mount preparation of the optic tectum in unlesioned (top) and at 24 hours post lesioned (bottom) Tg(mpeg1.1:eGFP) animals. Scale bars B: C:. TeO: tectum optic; SM: Stratum Marginale; SO: Stratum Opticum; SFGC: Stratum Fibrosum et Greseum Superficale; SAC: Stratum Griseum Centrale; SPV: Stratum Album Centrale; ON: optic nerve; R: rostral; C: caudal.

### 3.4. Following photoreceptor injury-induced chemotaxis, phagocytosis, and proliferation, microglia return their original niches

One of the hallmarks of activated microglia is their ability of phagocytosis and proliferation. In the subretinal space, individual microglia display enlarged somata and thickened processes that terminate in phagocytic cups (Fig. 4A). Spatial and temporal dynamics of microglial proliferation using immunostaining against proliferation marker, PCNA, following photoreceptor injury showed that at 24 hpl, 6.8% of 4C4+/Hoechst+ microglia were stained for PCNA (n=88 microglia from 6 animals). At 48 hpl the proportion of PCNA+ microglia increased to 59.4% (n=288 microglia from 6 animals) (Fig. 4B). Among the PCNA+ microglia, 97.1% were in the subretinal space. The peak of proliferation at 48 hpl suggests that signals that induce microglial migration are distinct from those that stimulate proliferation. Resolution of inflammation is critical for restoring tissue homeostasis. It requires clearance of cellular debris, cessation of the expression of pro-inflammatory molecules, and production of pro-resolving mediators, all which are associated with phenotypic switch of microglia from pro-inflammatory to anti-inflammatory states. At 3 dpl, microglia located in the subretinal space begin to migrate to their original locations among the somata of the RPE and laminae within the retinal parenchyma (Fig. 4C,D). Migrating parenchymal microglia adopt radial orientations with thin processes spanning the nuclear layers (Fig. 4E), and we infer this to be evidence of microglia returning to their respective niches in the inner retina. At 5 dpl, parenchymal microglia reacquire ramified morphologies, approximating that in unlesioned retinas (Fig. 4F,G). At 14 dpl, the number and morphologies of both parenchymal and sub-retinal microglia resemble those in unlesioned retina (Fig.4H,I). Together, these observations reveal a rapid but transient microglial response to photoreceptor injury. Microglial migration begins within hours of injury, proliferation peaks within 2 days, and microglial distribution and morphology largely return to their unlesioned state by 14 dpl. This timeline parallels the progression of photoreceptor injury and regeneration: extensive photoreceptor cell death occurs during the first few days after lesion, followed by regenerative neurogenesis and progressive restoration of the photoreceptor population. Thus, microglial redistribution and proliferation occur predominantly during the early injury and proliferative phases and largely subside as retinal regeneration approaches completion, suggesting that microglial responses are tightly coupled in time to the regenerative process.

**Figure 4.**
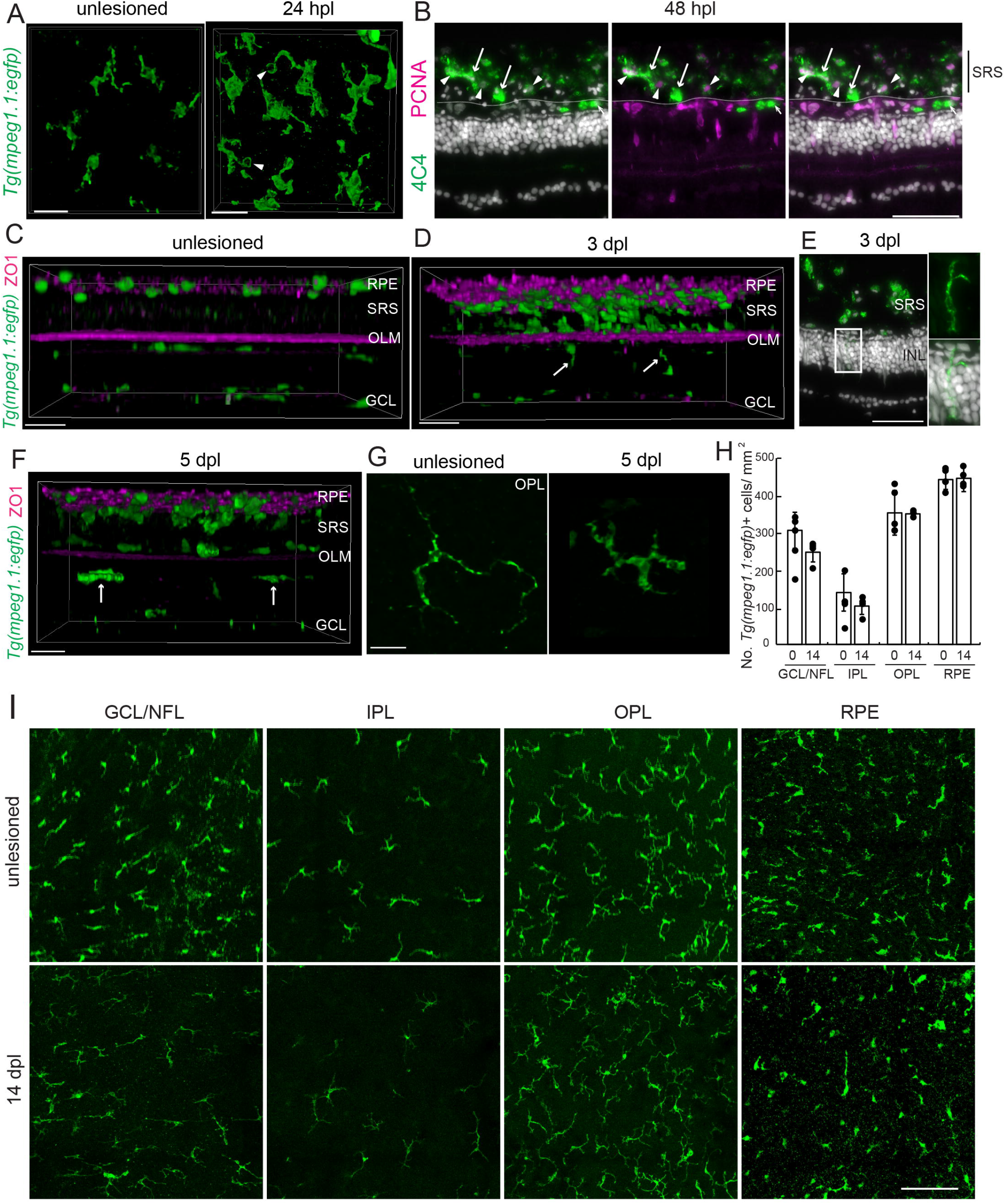
Following photoreceptor injury-induced chemotaxis, phagocytosis, and proliferation, microglia re-occupy their original niches. (A) 3D rendering of Tg(mpeg1.1:egfp)-labeled microglia in the subretinal space at 24 hpl. Arrowheads indicate phagocytic cups formed at the terminal of lateral processes. (B) Retinal cross section immunestained for 4C4 and PCNA at 2 days post lesion. Arrowheads and arrows indicate 4C4+/Hoechst+/PCNA+ proliferating microglia and 4C4+/Hoechst+/PCNA− non-proliferating microglia, respectively. Dotted lines indicate the outer limiting membrane. (C,D) Orthogonal 3D rendering of retinal flat-mount Z-stack series of Tg(mpeg1.1:eGFP) retina immunostained for ZO1 in unlesioned (C) and at 24 hours post lesioned (D) samples. Arrows indicate radially orienting microglia in the retinal parenchyma. (E) Retina cross section immunolabeled for 4C4 at 3 days post lesion. (F) Orthogonal 3D rendering of retinal flat-mount Z-stack series of Tg(mpeg1.1:eGFP) retina immunostained for ZO1 at 5 days post lesion. Arrows indicate microglia in the outer plexiform layer. (G) High magnification image of microglia in the outer plexiform layer of unlesioned and 5 days post lesioned retinas. (H) Quantification of the number of microglia in different retinal layers of unlesioned and 14 days post lesioned temporal retinas. (I) Maximum projection images of flat-mount retinas from the Tg(mpeg1.1:egfp) line in different retinal layers in unlesioned (top) and 14 days post lesioned (bottom) samples. Scale bars: A, C, D, E, F, G: 20um; B: 50um; I: 100um. RPE, retinal pigment epithelium; SRS, subretinal space; ONL outer nuclear layer; OPL outer plexiform layer; INL, inner nuclear layer; GCL/NFL, ganglion cell layer/nerve fiber layer.

### 3.5. Csf1ra is required to establish or maintain parenchymal microglia, but not subretinal microglia

Colony stimulating factor 1 receptor is a tyrosine kinase receptor that is essential for development, maintenance, and homeostasis of microglia. During development in mice, genetic mutation of Csf1r results in nearly complete absence of microglia(Dai et al. 2002; Erblich et al. 2011; Ginhoux et al. 2010). Zebrafish possess two homologs of the mammalian CSF1R gene, *csf1ra* and *csf1rb*. With the loss of functional mutation of *csf1ra*, invasion of early macrophages into the nervous system is compromised(Herbomel et al. 2001), and in adults the number of microglia is significantly less than in wildtypes(Oosterhof et al. 2018). We sought to characterize laminar distribution of retinal microglia in the *csf1ra* mutants and determine if following photoreceptor injury the absence of Csf1ra alters the chemotactic and/or proliferative response of microglia. For these experiments, we crossed the Tg(mpeg1.1:mCherry) reporter into the csf1ra^j4e1^ mutant. (For unknown reasons, csf1ra^j4e1^ mutants carrying mpeg1.1:egfp construct resulted in markedly reduced fertility.) In this line, the number of parenchymal microglia in the *csf1ra* mutants was dramatically less than in wildtype animals, and there was nearly a complete absence of microglia at the interface of the inner nuclear and inner plexiform layers (Fig. 5A,B). In marked contrast, the number of subretinal microglia was comparable between wildtypes and mutants (Fig. 5A,B). These observations indicate that signaling through Csfr1a is required either to establish the initial colonization or maintain parenchymal microglia. Further, these data show that the colonization/maintenance of microglia in the subretinal space is independent of Csf1ra. We next asked whether the anatomically distinct parenchymal and subretinal microglial populations could be distinguished transcriptionally based on expression of the Csf1r receptors. Both *csf1ra* and *csf1rb* were expressed across the microglial clusters, with no apparent enrichment of *csf1ra* in a specific population(Fig. 1F). Thus, the transcriptional profiles obtained from unlesioned retinas did not resolve the anatomically distinct microglial populations based on Csf1r expression. We next asked whether loss of Csf1ra alters the microglial response to photoreceptor injury. We first examined if the mutation in csf1ra alters death of photoreceptors following photolytic lesion. Comparison of the number of TUNEL-positive cells found no difference between wildtype and the Csf1ra mutants at 1 dpl (wildtype 13.95±0.53 cells, mut 14.45±2.22 cells) and 2 dpl (wildtype 8.43±2.07 cells, mut 6.90±2.04 cells). In the Csf1ra mutants at 24 hpl, activated microglia were present in the subretinal space, and microglia in the retinal parenchyma are largely depleted (Fig. 5C,D), consistent with the migration of microglia observed in wildtype retinas. In contrast, in the Csf1ra mutants there was significantly less microglial proliferation (Fig. 5F). At 2 and 3 dpl, 4C4-positive, PCNA-labeled microglia were only rarely observed (2 dpl, 0 PCNA+ microglia in total 45 microglial cells, 4 animals: 3 dpl, 2 PCNA+ microglia in total 44 microglial cells, 4 animals). These results indicate that Csf1r signaling is required for proliferation among activated microglia. Importantly, Müller glial proliferation and the generation of Müller glia-derived progenitors occurred normally in *csf1ra* mutants, and photoreceptor regeneration proceeded comparably to wildtype animals (data not shown), consistent with previous findings that photoreceptor regeneration proceeds normally in other microglia-deficient zebrafish mutants(Song et al. 2024). Thus, despite impaired microglial proliferation, loss of Csf1ra does not substantially disrupt the regenerative response of the retina. Finally, we examined the termination of the microglial chemotactic response in Csf1ra mutants. At 14 dpl, the number of microglia in the subretinal space of *csf1ra* mutants was significantly less than in the unlesioned mutant retinas (Fig. 5G,H). Concurrently, there were significantly more microglia within the outer plexiform layer (Fig.5G,I). These microglia display ramified, ‘resting’ morphologies, resembling microglia in unlesioned retinas (Fig. 5I). These results indicate that Csf1r signaling is not required for the acute migration of microglia toward injured photoreceptors but is required for the subsequent restoration of their homeostatic distribution.

**Figure 5.**
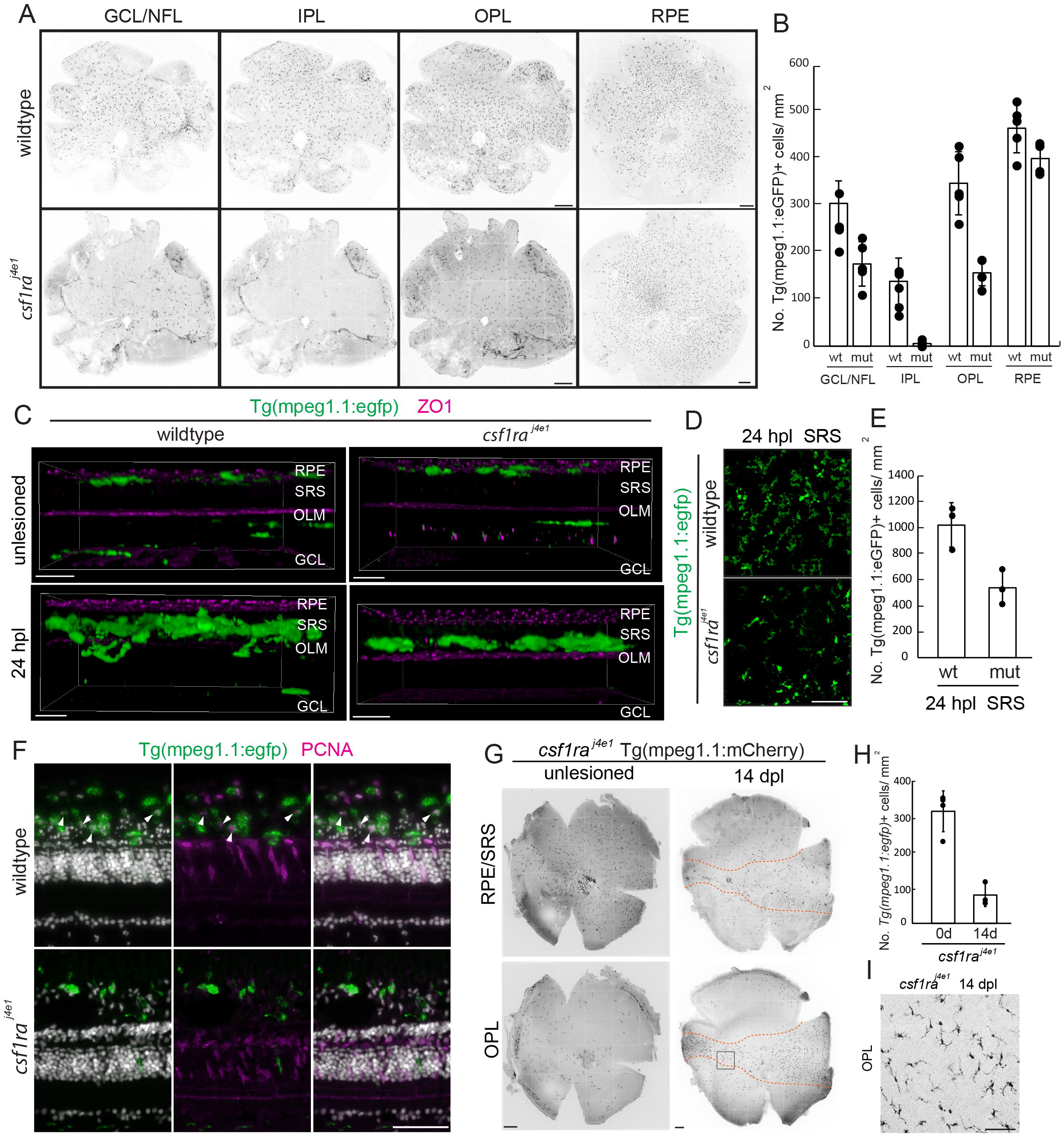
In the Csf1ra mutant microglia respond to photoreceptor injury normally by chemotaxis. but proliferation and redistribution during inflammatory resolution is compromised. (A) Maximum projection images of flat-mount retinas from the Tg(mpeg1.1:egfp) (top) and csf1ra^j4e1^; Tg(mpeg1.1:mCherry) lines in different retinal layers in unlesioned samples. (B) Quantification of the number of microglia in different retinal layers of wildtype and csf1ra mutant retinas. (C) Orthogonal 3D rendering of retinal flat-mount Z-stack series from microglia reporter line in wildtype and csf1ra^j4e1^ immunostained for ZO1 in unlesioned and at 24 hours post lesioned samples. (D) Maximum projection of microglia in the subretinal space in unlesioned and 24 hours post lesioned csf1ra^j4e1^ mutant animals. (E) Quantification of the number of microglia in the subretinal space in wildtype and csf1ra^j4e1^ mutant animals at 24 hours post lesion. (F) Cross section of wildtype (top) and Csf1ra mutant (bottom) retinas immunolabeled for 4C4. Arrowheads indicate 4C4+/Hoechst+/PCNA+ proliferating microglia. (G) Maximum projection images of flat-mount retinas from the Tg(mpeg1.1:egfp) (top) and csf1ra^j4e1^; Tg(mpeg1.1:mCherry) lines in different retinal layers in unlesioned and 14 days post lesioned retinas. (H) Quantification of the number of microglia in the subretinal space in csf1ra^j4e1^ mutant animals in unlesioned and at 14 days post lesion. (I) High magnification image of microglia in boxed region shown in (G). Scale bars: A, G: 200um; C, D: 20um; F: 50um. RPE, retinal pigment epithelium; SRS, subretinal space; ONL outer nuclear layer; OPL outer plexiform layer; INL, inner nuclear layer; GCL/NFL, ganglion cell layer/nerve fiber layer.

## 4. Discussion

Microglial heterogeneity has been increasingly recognized across the CNS(Fumagalli et al. 2025), with single-cell studies revealing distinct transcriptional states that vary according to location(Masuda et al. 2019), developmental stage(Hammond et al. 2019), and disease state(Fumagalli et al. 2025)(Keren-Shaul et al. 2017). In the zebrafish retina, following retinal damage microglia undergo dynamic transcriptional changes, including proliferative and inflammatory states(Mitchell et al. 2019);(Xu et al. 2025; Weimar et al. 2026; Ravishankar et al. 2026; Mitra et al. 2026). Our identification of proliferative, Cq1-enriched, and ApoC1/Apoeb-enriched microglial populations in the absence of retinal injury suggests that substantial microglial heterogeneity is present in the zebrafish retina. Although we cannot determine whether these transcriptional states correspond to specific anatomical niches, their presence in the unlesioned retina raises the possibility that microglial heterogeneity in the retina reflects the interaction between cellular state and local tissue environment. It remains unclear if the transcriptional diversity observed following retinal injury builds upon pre-existing heterogeneity among resident microglia. Although subretinal microglia is not present in the mammalian retina, multiple lines of evidence support the identification of the cells we observed in the subretinal space of zebrafish as *bona fide* microglia. The mpeg1.1:egfp-positive cells located among the somata of RPE consistently express the validated microglia marker, 4C4. Tg(mpeg1.1:egfp)+ cells among RPE form a regularly spaced, tangential mosaic, resembling the parenchymal microglia. Following photoreceptor injury, the mpeg1.1:egfp-positive cells among the RPE rapidly adopt activated morphologies, migrate into the photoreceptor layer, display phagocytic cups and appear to phagocytose photoreceptor somata and debris, hallmarks of the injury-induced responses of microglia observed in all vertebrate retinas(Silverman and Wong 2018; Rashid et al. 2019; Guo et al. 2022).

An outstanding, unanswered question raised by our observations is the homeostatic function of subretinal microglia. In homeostatic conditions, microglia play supportive and surveillance roles of the microenvironment, participating in synaptic pruning and plasticity, phagocytosing cellular debris, and maintaining the blood-brain barrier(Li and Barres 2018; Schafer and Stevens 2015; Gullotta et al. 2023). Based on their location, we infer that the subretinal microglia provide a form of immune surveillance confined to the subretinal space and, therefore, provide exclusive service to photoreceptors and/or RPE. One may speculate that subretinal microglia may phagocytose photoreceptor outer segments or mediate other aspects of homeostasis in the subretinal space, such as maintaining choroid-retinal barrier formed at the RPE layer. We propose that in a healthy retina subretinal microglia cooperate closely with RPE cells to achieve these tasks. Our observations also raise the question of why subretinal microglia are not present in mammalian retinas. Determining the developmental origin and ontogeny of subretinal microglia in zebrafish may provide insight into how microglial localization, specialization, and interaction with retinal cells have evolved across vertebrates.

Despite their transcriptional and positional heterogeneity, microglia across the retina exhibit a highly conserved response to acute photoreceptor injury. Following photolytic damage, microglia rapidly migrate toward and accumulate within the subretinal space, followed by proliferation and gradual return toward homeostatic state. These results suggest that microglial heterogeneity does not necessarily predict fundamentally different responses to acute tissue damage. The photolytic lesion paradigm used in this study provides several advantages for resolving the spatial and temporal dynamics of this response. First, unlike lesion paradigms that disrupt blood vessels(Xu et al. 2025), a photolytic lesion in the avascular retina of zebrafish does not involve infiltration of peripheral circulating immune cells. This reduces the cellular complexity of the inflammatory response and allows investigation of inflammatory events that can be largely ascribed to resident microglia. Second, in contrast to wide-spread neuronal death induced by neurotoxins(Mitchell et al. 2018), photolytic lesions result in the selective death of photoreceptors in a manner that is spatially restricted. This allows comparisons of microglia that reside within and outside the region of damaged photoreceptors. Third, using newly developed flat-mount retina/RPE preparations from multiple timepoints covering both the acute and resolution phases of inflammation, we were able to qualitatively and quantitatively determine the spatial and temporal responses of microglial activation and recovery following a photoreceptor injury. Together, these features allowed us to quantitatively characterize the spatial and temporal dynamics of microglial migration, accumulation, proliferation, and recovery following photoreceptor injury. In addition to defining the general dynamics of microglia, we describe a unique population of microglia, not present in mammals, that resides in the subretinal space among the somata of the RPE. To our knowledge, this is the first formal characterization of this subretinal microglial population and their responses following photoreceptor injury.

Following photoreceptor injury, nearly all parenchymal microglia underlying the injured photoreceptors rapidly migrate radially into the subretinal space. This response parallels observations in mammalian models of photoreceptor degeneration, in which retinal microglia relocate from their homeostatic niches to the outer retina and subretinal space following photoreceptor injury(Santos et al. 2010; Joly et al. 2009; O’Koren et al. 2019). In mice, microglia from distinct retinal niches relocate to the subretinal space following degeneration, demonstrating that injury can override their normal anatomical organization. Our findings extend these observations to zebrafish and suggest that parenchymal microglia in the zebrafish retina share conserved injury-responsive properties with their mammalian counterparts. The extensive radial reorganization of parenchymal microglia also indicates that photoreceptor injury generates signals capable of influencing microglia well beyond the immediate site of photoreceptor death. Rather than remaining confined to their homeostatic positions, microglia throughout the retinal depth are recruited toward the injured outer retina, suggesting that injury-associated signals propagate across retinal layers and coordinate a tissue-wide microglial response. The nature of these signals remains to be determined and could involve direct signals from damaged photoreceptors as well as secondary signals from neighboring retinal cells. Notably, microglial activation in the optic tectum was negligible following retinal injury, arguing against a generalized or systemic activation of microglia and instead supporting the predominantly local nature of the retinal response. Together, these findings suggest that photoreceptor injury induces a highly coordinated, retina-restricted reorganization of resident microglia, in which anatomically distinct microglial populations converge on the injured outer retina.

In response to photoreceptor injury, microglia rapidly alter their morphologies, from ramified to rod-shaped, bushy, and amoeboid shapes. Distinct microglial morphologies reflect their different function(Walker et al. 2014; Holloway et al. 2019). The immediate and transient appearance of rod-shaped microglia suggests rapid migration following tissue damage. Consistent with this, in vivo time-lapse images of microglia following laser-induced neuronal injury in zebrafish larvae demonstrated microglia rapidly polarize their cellular processes before migrating toward the lesion site(Sieger et al. 2012; Sieger and Peri 2013). The elongated and highly polarized morphology of rod-shaped microglia may therefore represent a transient morphological state that facilitates rapid migration toward the site of injury. Following the onset of the photolytic injury, microglia can initiate chemotaxis within 60 minutes. We speculate that these rapid morphological transitions may be accompanied by changes in microglial gene expression. Importantly, the rapid emergence of activated microglial phenotypes precedes the appearance of TUNEL-positive, pyknotic nuclei in the outer nuclear layer. This temporal relationship is consistent with the possibility that microglia respond to early injury associated with signals before extensive photoreceptor apoptosis becomes morphologically evident. At the injury site, microglia subsequently engage in active phagocytosis of cellular debris generated by dying neurons(Mitchell et al. 2018). Efficient clearance of dying cells contributes to the subsequent transition from a pro-inflammatory toward anti-inflammatory state. Together, our observations suggest that microglial responses to photoreceptor injury are temporally coordinated.

Our results show that microglial proliferation is spatially and temporally controlled. The delay between chemotaxis and proliferation suggests that these processes are governed by distinct mechanisms. We identified Csf1 receptor signaling as a critical regulator for microglial proliferation during acute inflammation, and this is consistent with studies in mammals, where CSF1R ligands, CSF1 or Il34, stimulate microglial proliferation(Yamamoto et al. 2010; Mizuno et al. 2011), whereas pharmacological inhibition of CSF1 R suppresses proliferation following injury or during neurodegeneration(Gómez-Nicola et al. 2013; Gerber et al. 2018). Identifying the cellular sources of Csf1 ligands will be an important next step toward understanding the cell-cell interactions that regulate this response.

At 14 dpl, *csf1ra* mutants contained significantly fewer subretinal microglia but more parenchymal microglia than in the retinas of unlesioned mutants. Because Csf1ra mutants have impaired microglial proliferation, the increase in parenchymal microglia is unlikely to result from local proliferation. One possibility is that microglia are recruited from outside the retina, such as through the optic nerve, as elongated microglia are frequently observed along optic nerve axon bundles. Alternatively, activated subretinal microglia may fail to return to their original location and instead colonize the retinal parenchyma, particularly the outer plexiform layer. We propose that Csf1r signaling may be required for the repositioning of subretinal microglia during inflammatory resolution. Recent scRNA-seq data show that the Csf1 ligand Il34 is strongly induced in the RPE following genetic RPE ablation(Leach et al. 2021), suggesting that IL34–CSF1R signaling between RPE and subretinal microglia may promote their recolonization of the subretinal space. These findings suggest that distinct molecular mechanisms may regulate microglial colonization of the retina during development and the resolution of inflammation.

## 7. Author Contributions

MN: Conceptualization, Writing-review and editing, Data curation, Formal Analysis, Investigation, Methodology, Validation, Visualization, Writing-original draft. LJ, SG: Data curation, Formal Analysis, Investigation, Methodology, Validation, Visualization. PH, TH: Conceptualization, Writing-review and editing, Funding acquisition, Project Administration, Resources, Supervision.

## 9 Acknowledgments

We thank Mr. Dilip Pawer and Mr. Garrison Ader for zebrafish care and husbandry.

## 10 Data availability statement

The original contributions presented in the study are included in the article, further inquiries can be directed to the corresponding author.

